# Isolation and characterization of TayeBlu, a novel bacteriophage of *Azotobacter vinelandii*

**DOI:** 10.1101/2025.04.23.650044

**Authors:** Taiwo Mercy Akanbi, Mariana Labat, Tyler Sun, Derek A. Smith, Sarah C. Bagby

## Abstract

Soil microbial communities drive global biogeochemical cycles and alter crop yields through nitrogen fixation. As agents of genetic mobility, mortality, and nutrient release, viruses have been shown to influence microbial community structure and activity in numerous marine and aquatic systems. However, their impacts on terrestrial ecosystems are less well understood, in part because few model phage-host systems have been established for soils. To fill this gap, we sought to develop a new model system for viral infection of nitrogen-fixing bacteria derived from agricultural soil. Here, we report the isolation, characterization, and sequencing of the novel bacteriophage TayeBlu, which infects the globally distributed aerobic soil bacterium *Azotobacter vinelandii*, a facultative diazotroph. TayeBlu was isolated from the rhizosphere of tomato plants in a farm greenhouse. We find that the availability of nitrogen to host cells strongly influences TayeBlu infection physiology at the level of adsorption kinetics, time to lysis, and burst size. Taxonomic and comparative genome analysis reveal that TayeBlu belongs to an understudied family in class *Caudoviricetes* in which a small core of structural and assembly genes has persisted through adaptive diversification on different bacterial hosts.

**IMPORTANCE:** Understanding the forces regulating soil microbial activity is critical for building accurate ecosystem models that can inform land-management strategies aimed at mitigating climate risks and stabilizing the global food supply. For agricultural sustainability, it is particularly important to understand the dynamics of soil nitrogen-fixing bacteria like *Azotobacter vinelandii*, a well-studied and globally distributed species whose activity promotes plant growth and soil fertility. To support detailed investigations of the impact of viruses on diazotroph ecosystem outputs, we isolated and investigated a novel soil virus that infects *A. vinelandii*. This new phage-host system holds promise as a model experimental system for soil viral studies, illuminating a critical but poorly understood aspect of soil ecology.

## INTRODUCTION

Soil microbial communities shape global biogeochemical cycles and local ecosystem function through metabolic processes and ecological interactions that determine the bioavailability of carbon (C) and nitrogen (N) [1, 2], affecting plant productivity and soil function. Within these communities, N-fixing bacteria (diazotrophs) play a crucial role by converting atmospheric N_2_ into biologically available forms. Annually, biological N fixation contributes ∼200M metric tons of bioavailable N to Earth’s ecosystems [3, 4], including nearly half the total N present in crop fields [5, 6]. By releasing organic N as a public good, soil diazotrophs facilitate the growth of plants and other microbes [7, 8], which otherwise may scavenge N by degrading recalcitrant soil organic matter [7, 8], potentially increasing carbon dioxide emissions [9].

Infection of bacteria by their viruses (bacteriophages, or phages) alters host ecological impacts. Phages influence nutrient cycling and control bacterial population dynamics and mortality [10]. In addition to direct lysis, phage infection impacts bacterial communities and biogeochemical processes through several distinct mechanisms: phages can reprogram host metabolism by introducing auxiliary metabolic genes (AMGs) [11], accelerate nutrient turnover through release of host cellular contents [12], and transfer genetic material between hosts [13, 14]. Phage infection dynamics can be highly dependent on host physiological state, with low-nutrient conditions reducing phage infectivity and burst size (e.g., [15, 16]). Viral lysis accounts for substantial bacterial mortality and subsequent nutrient release in well-studied marine systems [12, 14, 17]. While available evidence suggests that soil environments contain substantial phage populations (∼ 10^3^–10^9^ viral particles per gram of soil depending on soil type and conditions [18]), our knowledge of phage-host dynamics in soil ecosystems lags considerably behind marine systems [19, 20].

A major challenge in soil virology is the limited number of established phage-host systems available for experimental investigation, with current models capturing only a handful of the predominant soil bacterial phyla [21, 22]. This limitation is especially pronounced for N-fixing bacteria, a problematic gap given the critical importance of diazotrophy for soil function. In particular, the observation that N flow to the soil system can differ substantially between diazotrophs (e.g., between plant-associated and free-living species) [23] points to a need for diverse experimental systems to support comparative analysis of the ecosystem impacts of phage infection of different diazotrophic hosts. Recent work has identified phages infecting *Klebsiella* sp. M5al [24], a facultative diazotroph from a genus that often grows in association with a plant host. *Klebsiella* spp. fix N anaerobically, with low levels of O_2_ rapidly inhibiting nitrogenase, the enzyme responsible for N fixation [25]. By contrast, the globally distributed soil bacterium *Azotobacter vinelandii* is a free-living and obligately aerobic facultative diazotroph [26, 27]. Azotobacters are noteworthy for their specialized metabolic adaptations (e.g., extraordinarily high respiration rates) to protect nitrogenase from oxygen (reviewed in [26]), offering an opportunity to investigate infection physiology under widely different host metabolic states. With a versatile metabolism, a long history as a soil model system, and modern tools for mechanistic investigation [28, 29], *A. vinelandii* is an ideal host for phage model system development. Despite numerous reports of characterized *A. vinelandii* phage isolates several decades ago [30, 31, 32], to our knowledge none of these phages remain in cultivation, and none have been sequenced or characterized by modern methods.

To fill this gap, we sought to isolate novel phage of *A. vinelandii* that can be developed as an experimental model system for investigation of phage impacts on free-living soil diazotrophs. Here, we report the isolation, characterization, and sequencing of the novel rhizosphere siphovirus TayeBlu infecting *A. vinelandii* strain OP, and we demonstrate that phage infection is significantly influenced by N availability in the medium. Through comprehensive genomic analysis, we identify this phage as belonging to a novel viral family within an unclassified order of the class *Caudoviricetes*, with a conserved core of structural and replication genes but low genomic similarity to other known soil phage. This novel phage shows promise as an experimental system for studies on the influence of soil phages on N-fixing bacterial communities in variable soil environments and their impact on global biogeochemical cycles [19].

## MATERIALS AND METHODS

### Host strain, growth media, and phage buffer

All experiments were performed with *Azotobacter vinelandii* str. OP (hereafter *A. vinelandii*), which was the generous gift of Xinning Zhang. *A. vinelandii* was grown in three media spanning high and low C and N availability. PYCa (Peptone-Yeast-Calcium) is a rich and N-replete medium consisting of 15 g L^-1^ peptone, 1 g L^-1^ yeast extract, 1g L^-1^ dextrose, and 4.5 mM CaCl_2_. Dean’s Burke (Dean’s B) medium is a minimal, N-free medium consisting of 1 mM phosphate buffer, 20 g L^-1^ sucrose, 200 mg L^-1^ MgSO_4_, 91.1 mg L^-1^ CaCl_2_, 1 µM Na_2_MoO_4_, and 5.1 mg L^-1^ FeSO_4_. As an intermediate condition, we supplemented Dean’s B with ammonium chloride (1 mM NH_4_Cl final concentration) as an N source (AC_Dean). Phage buffer consisted of 68 mM NaCl, 1 mM CaCl_2_, 10 mM MgSO_4_, and 10 mM Tris, pH 7.5.

### Isolation and purification of phage TayeBlu

Phage TayeBlu (named from "Taye", a Nigerian-Yoruba name for "first child to taste the world," reflecting its status as the first *A. vinelandii* phage isolate to be characterized through modern methods, and "Blu", a local reference reflecting the isolate’s Cleveland origin) was isolated from rhizosphere soil collected at the base of a tomato plant in a greenhouse at the Case Western Reserve University Farm, Hunting Valley, Ohio (41^◦^29^′^36*^′′^*N, 81^◦^25^′^24*^′′^*W) on August 30, 2023. Using a hand shovel sterilized with 70% ethanol, 5055 grams of dry, dark soil were collected from within 2 cm of the plant base (Fig. S1), where fine roots were visible. Ambient temperature was 18 ^◦^C. Samples were stored on ice and kept at 4 ^◦^C until further processing.

Phage isolation used the enrichment method [33], followed by spot test analysis andagarose overlays [34]. Briefly, fresh soil samples (∼25 g) were enriched with 500 µL of an overnight *A. vinelandii* culture and incubated at 30 ^◦^C for 48 hours with shaking at 225 rpm. The mixture was centrifuged at 4700 × g for 30 min, and the supernatant was passed through a 0.22-µm PES filter to remove bacterial cells. Filtrates were screened for phage activity using the spot test method [35] with *A. vinelandii* cultured in the three media described above. Briefly, 500 µL of overnight culture was mixed with 4.5 mL of 0.6% molten agar and spread evenly on a 1.6% agar plate. After solidification of the top agar matrix, 10 µL of filtrate was spotted onto the surface and allowed to dry. Plates were incubated at 30 ^◦^C for 24 hours and monitored for plaque formation. The presence of a clear or turbid clearing on the plated bacterial lawn was considered a putative phage plaque.

Putative phage plaques were cored and suspended in 100 µL phage buffer, serially diluted, and then plated using the agarose overlay method; 10 µL of the target dilution (typically 10^-4^–10^-6^) was added to 500 µL overnight culture and 4.5 mL of 0.6% molten agar and spread evenly on a 1.6 % agar plate. Plates were incubated at 30 ^◦^C overnight. For three consecutive rounds of purification, a single, isolated plaque was resuspended in 100 µL of phage buffer, diluted, and plated to obtain a clonal phage population. Purified clonal phage was then amplified, titered, archived (4 ^◦^C for immediate use; *—*80 ^◦^C in 6.54 % DMSO or 20 % glycerol for freezer stock) and used for subsequent characterization in this study [35].

### Phage-host adsorption kinetics

To determine the adsorption rate constants (*k*) of phage TayeBlu and *A. vinelandii* strain OP in PYCa, Dean’s B, and AC_Dean media, we performed adsorption assays following previously described protocols [36, 37]. Briefly, in each medium, triplicate cultures of *A. vinelandii* were grown to mid-exponential phase (∼ 10^8^ colony-forming units (CFU) per mL) as determined by optical density measurements and OD-to-CFU calibration curves established previously for each medium. An aliquot of 2 *×* 10^8^ cells was removed from each culture and transferred to a 5-mL glass culture tube and the volume was adjusted to 2 mL with fresh medium, to achieve a culture density of 1 *×* 10^8^ CFU mL^-1^. High-titer TayeBlu lysate (4.3–7.3 *×* 10^10^ PFU mL^-1^) was added to a multiplicity of infection (MOI) of 0.1.

Immediately following infection (*t*_0_) and at predetermined intervals, samples were collected for both total and free phage quantification. For free phage measurements, samples were immediately filtered through 0.22-µm PES filters to remove bacterial cells and any cell-associated phages. For total phage measurements, parallel samples were left unfiltered. All samples were serially diluted in phage buffer and enumerated at two sequential dilutions using the agarose overlay method. AC_Dean was used as the base medium for plaque assays from AC_Dean adsorption experiments; PYCa was used for plaque assays from experiments in both PYCa and Dean’s B. This use of PYCa ensured reliable plaque formation, as our preliminary experiments demonstrated that PYCa provides consistently good conditions for TayeBlu plaque formation on this host. Plates were incubated at 30 ^◦^C and plaques were enumerated after 24, 48 and 72 hours to ensure complete development of all viable plaques.

Statistical analyses of the decrease in free phage concentration over time were performed using R (version 4.4.3) [38] with the tidyverse [39], Hmisc [40], errors [41], ggtext[42], BSDA [43], and patchwork [44] packages. Data were filtered to exclude dilutions with zero counts and or plaques too numerous to count (TNTC). For each measurement, the phage concentration (plaque-forming units (PFU) per mL) was calculated and assigned an uncertainty scaled to the square root of the number of plaques observed (truncated at a minimum of 1.2), reflecting Poisson error in counting statistics. Weighted means were calculated for each time point, with weights assigned as the inverse of the relative error for each measurement. Following [36], the fraction of free phage remaining (*P_t_*/*P*_0_) was plotted against time. Adsorption rate constants (*k*, in units of mL/min) were determined by fitting the natural log-transformed data to a first-order kinetic model, *k* = *—* ln(*P_t_*/*P*_0_)/*Nt*, where *P_t_* is the concentration of free phage at time *t*, *P*_0_ is the initial free phage concentration, and *N* is the bacterial concentration. All error propagation used the errors package. Differences in fitted *k* values were tested with Welch’s modified two-sample t-test (BSDA tsum.test).

### One-step growth curves

We performed one-step growth experiments following previously described protocols [36, 45, 46]. Based on fitted adsorption rate constants (*k*), we chose media-specific experimental parameters to achieve tightly synchronized infection of a large fraction of the host population: for PYCa, a target MOI of 3 and a 3-minute adsorption period; for AC_Dean, a target MOI of 4 and a 7-minute adsorption; and for Dean’s B, a target MOI of 6 and an 8-minute adsorption. In each one-step assay, triplicate cultures were grown in the test medium to mid-logarithmic phase (1–1.8 *×* 10^8^ CFU mL^-1^) as determined by optical density measurements and OD-to-CFU calibration curves established previously for each medium. Aliquots of 3.2 *×* 10^9^ cells were removed from each culture and transferred to an experimental and a control 250-mL flask, then adjusted to concentrations of ∼1 *×* 10^8^ CFU mL^-1^ using fresh medium. Pre-infection samples (2 mL total) were collected for archiving and OD measurement. High-titer TayeBlu lysate (4.3–4.5 *×* 10^10^ PFU mL^-1^) was added to achieve the target MOI in experimental flasks; parallel no-infection controls were amended with an equivalent volume of phage buffer. Working volumes during the adsorption period were 30 mL for Dean’s B and 31 mL for AC_Dean and PYCa.

After the adsorption period, to prevent further infection, 25 mL of each experimental and control culture was transferred to fresh 500-mL flasks and diluted tenfold by addition of 225 mL of fresh medium. Flasks were incubated at 30 ^◦^C with shaking at 225 rpm. At predetermined intervals, samples were collected for total phage (10 µL), free phage (10 µL filtered through 0.22-µm PES filters), and OD measurement. Phage abundance in free and total phage samples was enumerated using plaque assays on PYCa (PYCa and Dean’s B experiments) or AC_Dean (AC_Dean experiment). Plates were incubated at 30 ^◦^C for 24 hours in PYCa medium, or 48 hours in AC_Dean medium to allow for slower host growth in this medium. Plaque counts were stable after the incubation period. Plates were permitted to develop at room temperature for up to 24 hrs after incubation for additional re-examination, to facilitate accurate counting of the very small plaques obtained in AC_Dean. Analysis of plaque counts, assignment of uncertainties, and error propagation were conducted as described for phage-host adsorption kinetics. Burst size was calculated as the ratio of final free phage concentration to the initial number of infected cells, where the number of infected cells was determined by subtracting the initial free phage concentration from the initial total phage concentration. (Plaque counts in total phage measurements are the sum of plaques due to free phage and plaques due to infected cells; the latter are expected to yield one plaque per infected cell.) Differences in burst sizes were tested with Welch’s modified two-sample t-test (BSDA tsum.test). Latent period was determined as the time between phage adsorption and the onset of the release of phage progeny [36].

### Morphological characterization with transmission electron microscopy

Phage morphology was examined by transmission electron microscopy (TEM) following a protocol modified from [47]. Purified high-titer phage lysate (1 mL at 1 *×* 10^10^ PFU mL^-1^) was centrifuged at 20,630 × g for 50 min. The supernatant was carefully removed and replaced with 500 µL of 0.1 M ammonium acetate. The sample was centrifuged again at 20,630 × g for 50 min. For visualization, 5 *µ*L of the concentrated phage suspension was applied to a carbon/Formvar-coated copper grid (Electron Microscopy Sciences, FCF300CU50, purchased from Fisher Scientific, Catalog No. 50-260-36) and allowed to adsorb for 1 min. Without removing excess liquid, 5 µL of 2% uranyl acetate (w/v) was added for negative staining and left for an additional minute. Excess liquid was then wicked away using filter paper, and the grid was air-dried. Grids were examined using a TECNAI SPIRIT T12 TEM operating at 100 kV.

### DNA extraction and whole-genome Nanopore sequencing

High-molecular-weight DNA was extracted following a modified protocol based on [48] and the DNeasy PowerSoil Pro (QIAGEN, catalog no. 47014) DNA kit protocol. Briefly, phage lysate (6 mL) was treated with DNase I-RNase A (added at a ratio of 0.5 µL of 10 mg mL^-1^ stock per mL lysate, for a total of 3 µL nuclease mixture) and incubated at 37 ^◦^C for 30 minutes to eliminate contaminating nucleic acids. Phage particles were concentrated by adding 0.5 mL of 30% PEG-8000 per ml of treated lysate, followed by overnight incubation at 4 ^◦^C and centrifugation at 10,000 × g for 10 minutes. The resulting pellet was resuspended in 300 µL of 5 mM MgSO_4_ and transferred to PowerBead Pro tubes (QIAGEN, cat. no. 47014) for capsid disruption using an MP Biomedicals FastPrep 24 bead beater (6m s^-1^ for 30 s, repeated once). DNA purification continued according to the manufacturer’s protocol using the DNeasy PowerSoil Pro Kit (QIAGEN, catalog no. 47014). The extracted DNA was sequenced using the MinION 1KB long-read sequencer (Oxford Nanopore R10.4.1 flowcell) after library preparation with the SQK-NBD114.24 Native Barcoding Kit 24 V14 following the manufacturer’s guidelines.

### Phage genome assembly and annotation

Phage genomes were assembled using Oxford Nanopore Technology (ONT) sequencing data. Raw reads were basecalled using Dorado (v0.7.2) [49] with the super-accuracy model and 400 bps parameters [50]. Demultiplexing was performed with the –both-ends argument, followed by adapter trimming using the same software [49]. BAM files were converted to FASTQ format using Samtools (v1.21) [51]. Quality assessment of the FASTQ files was conducted using FastQC (v0.11.9) [52] and results were aggregated with MultiQC (v1.9) [53]. Quality trimming was implemented with Prowler [54] using the F1 option (Q-score threshold of 20) to retain the longest high-quality fragments. Assembly was performed using Metaflye (v2.9) [55] with the –meta option, which is optimized for viral genomes. The resulting assembly achieved an average read depth of >100x across the genome. Assembly graphs were visualized with Bandage [56] to confirm the presence of complete circular contigs. The completeness and potential contamination of the assembled viral genomes were assessed using CheckV [57].

Coding sequences were predicted using multiple tools including GeneMark, GeneMarkS, Glimmer (v3.02) [58], Prodigal (v2.6.3) [59], MetaGeneAnnotator (v1.0) [60], and phold (v0.2.0) [61]. Predictions were integrated using a weighted scoring system as described in [24], with scores calculated based on gene length, overlap, protein identification, and programming potential. Functional annotations were assigned based on significant alignment scores from BLASTP (e-value *<* 10^-5^) [62], Swiss-Prot (release 2023_01) [63], HHpred (probability *>* 90% and e-value *<* 1) [64], and phold [61], which converts protein sequences into 3Di structural token format for comparison against a repository of >1M phage protein structural models [65, 66, 67, 68, 69]. Phage genome visualization was performed using the final manually curated GenBank-formatted TayeBlu sequences and a modified version of the phold python plotting script (plot.py).

Assignments were manually curated to resolve conflicts between predictions with attention to genomic context; across tools, HHpred results were weighted most heavily and phold least, in line with current levels of benchmarking support for these tools. Additional genomic features, including tRNAs, introns, and spanins, were screened for using the structural and functional phage annotation pipelines in Apollo [70]. Rho-independent transcription terminators were identified using ARNold [71], which employs both RNAmotif and Erpin algorithms. Predicted terminators were manually curated based on their genomic context and structural features. Putative AMGs were screened using DRAM-v (v1.3.0) [72] with default parameters, retaining predictions with an M flag.

### Phage taxonomy classification

Identification of related phages and taxonomic classification followed a multi-tiered approach to ensure accuracy [73]. At the protein level, close relatives were recruited from Viral RefSeq database release 223 using vConTACT3 beta mode [74]. We examined the complete set of vConTACT3 taxonomic assignments, focusing on identifying the related viruses that cluster with TayeBlu at each taxonomic level and on the novel taxonomic groups assigned. We utilized ViPTree’s web interface to generate relatedness data [75]. At the nucleotide level, the complete genome of phage TayeBlu was compared against the NCBI non-redundant (nr) nucleotide database using BLASTN to identify the closest relatives. The average nucleotide identity (ANI) of the closest relative was estimated by multiplying the genome coverage by the percent identity of the hit. Phages identified by these approaches were submitted together with TayeBlu to VIRIDIC [76] for pairwise genomic distance analysis. VIRIDIC output was visualized by using a modified version of the original VIRIDIC R script, and the dendextend package [77] for dendrogram presentation.

For protein-based phylogeny, we employed vConTACT3 to assign taxonomy to TayeBlu using the NCBI Viral RefSeq database (release 223). The vConTACT3 algorithm constructs gene-sharing networks based on protein similarity and employs hierarchical clustering to assign taxonomy across multiple ranks from realm to genus level [74]. The analysis was performed using default parameters for protein clustering and network-based classification leveraging the International Committee on the Taxonomy of Viruses (ICTV) framework. Final taxonomic assignments were determined based on the optimal distance thresholds identified for each taxonomic rank. This analysis produced a hierarchical clustering of TayeBlu relative to known phages in RefSeq, GenBank, and specialized viral databases, identifying 42 viruses with significant similarity (SG values *≥* 0.020). We then applied the ICTV-recommended thresholds for taxonomic assignment (intergenomic similarity *≥* 95%, conspecific; *≥* 70–95%, separate species within a genus). The phylogenetic tree was visualized using the package ggtree (v3.10.1) in R (version 4.3.3) [78, 38].

### Comparative genome and core gene analysis

TayeBlu, the three best BLAST hits from NCBI nr, and the five other Viral RefSeq phages of novel_family_19 were analyzed for genome comparison and core gene identification. In order to perform these analyses, we downloaded the fasta files of all 8 phages from NCBI and performed genome annotation using pharokka with the -g prodigal-gv option [65, 79]. The resulting GenBank files, together with our manually curated TayeBlu .gbk file, were analysed and visualized for gene cluster synteny using clinker genome analysis with the default minimum alignment sequence identity (0.3) [80]. Core gene analysis was performed by clustering predicted protein sequences using MMseqs2 with a minimum sequence identity threshold of 30% and a coverage threshold of 80% (–min-seq-id 0.30 -c 0.8). The protein cluster results and Genbank annotations were further analyzed and visualized in R (version 4.4.3) with dplyr, tidyr, and stringr (from tidyverse v2.0.0) for data wrangling; and ggplot2 (v3.5.2), scales, and patchwork for plotting and multi-panel figure assembly [39, 44, 81, 38] for protein clustering distribution. Clusters with representatives from all 9 analyzed phages were considered core genes; those with representatives from 8 of the 9 were considered near-core.

### Comparison with PIGEON database environmental phages

We downloaded 145,535 metagenomic viral operational taxonomic units (vOTUs) from natural soil and rhizosphere environments from the PIGEON database (https://datadryad.org/stash/dataset/doi:10.25338/B8C934, accessed October 2024; all vOTUs from [82]). This dataset includes 53,391 vOTUs specifically derived from the tomato rhizosphere, an environment similar to the sample from which TayeBlu was isolated. Species-level clustering was performed using CheckV v0.8.1 and its associated scripts, anicalc.py and aniclust.py [57], with a threshold of 95% average nucleotide identity (ANI) over 80% genome coverage.

## RESULTS AND DISCUSSION

### Isolation and morphological characterization of novel phage TayeBlu

TayeBlu was successfully isolated from soil samples collected from the greenhouse farm enclosure at the CWRU University Farm (Hunting Valley, OH). As an initial investigation of the effects of nitrogen availability on infection, we asked whether plaque morphology differed across media. We consistently observed the clearest plaques on rich medium (PYCa; diameter ∼ 1.6 mm, Fig. **1**A) and tiny plaques on minimal medium amended with ammonium chloride as a nitrogen source (AC_Dean; ∼ 0.6 mm, Fig. **1**B, E). Plaques on minimal medium (Dean’s B) varied, with some plates showing temperate-like plaques (∼ 7.2 mm, Fig. **1**C) and others showing no clear plaques, though parallel inoculation of AC_Dean plates confirmed the presence of active phage in the inocula (Fig. **1**F; compare to panel E). Morphological characterization by transmission electron microscopy (TEM) revealed an icosahedral head with mean diameter 66.389 *±* 3.450 nm (*n* = 11) and a long, non-contractile tail (mean length 139.843 *±* 13.930 nm, *n* = 11; Fig. **1**D), features historically associated with siphoviruses [83].

**FIG 1.**
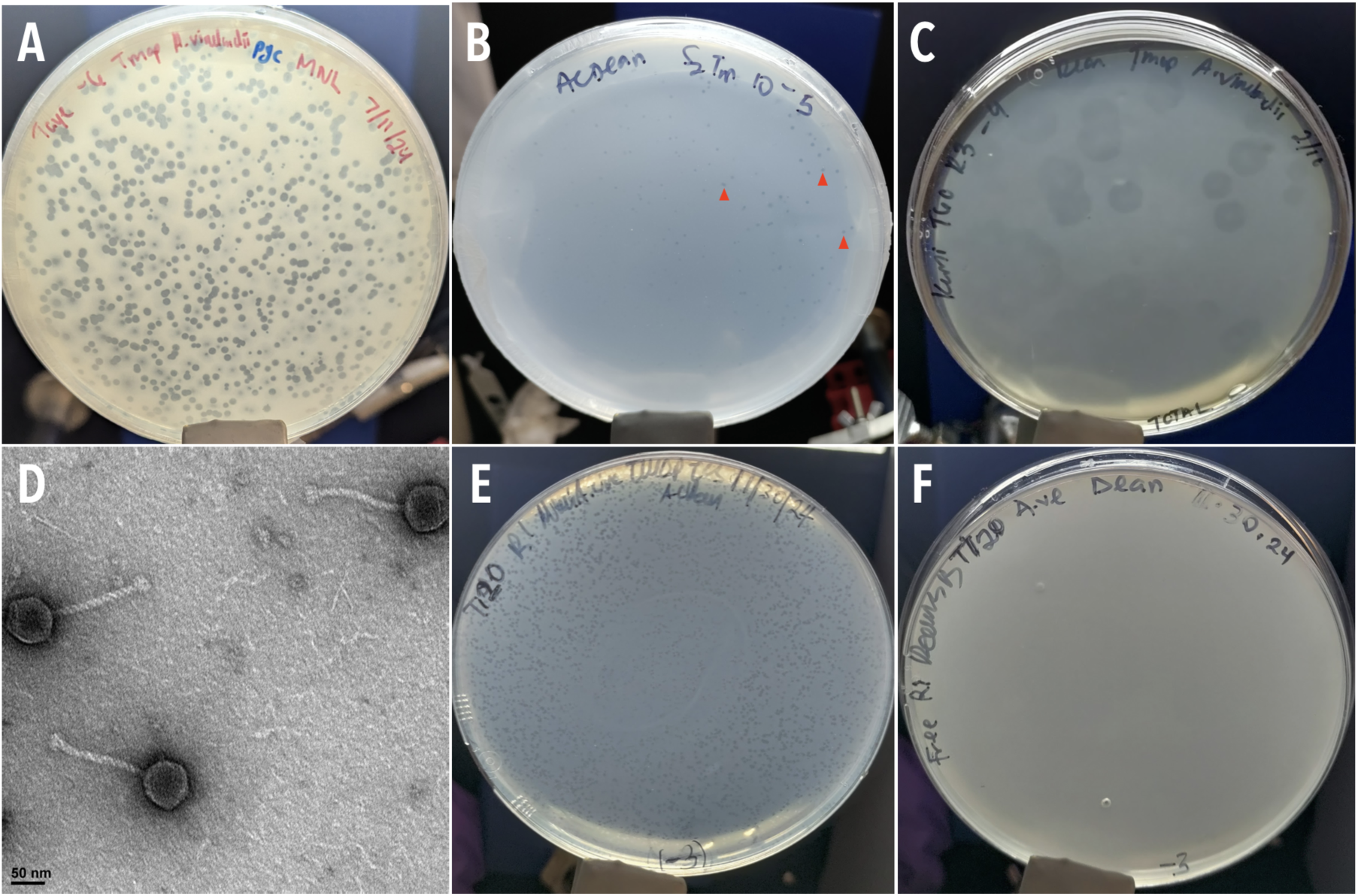
TayeBlu plaque and capsid morphology. (A) Plaques on *A. vinelandii* strain OP on rich medium (PYCa). (B, E) Plaques on N-replete minimal medium (AC_Dean). Red arrowheads in panel B mark representative plaques. (C, F) Large temperate-like plaques (C) and no discernible plaques on minimal medium (Dean). Plates in (E) and (F) were inoculated in parallel from the same phage stock. (D) Negative-stained transmission electron micrograph of phage particles.

### Infection dynamics

Phage TayeBlu infection dynamics on *A. vinelandii* strain OP in liquid medium, like its plaque morphologies on solid medium, were significantly influenced by media composition, with effects spanning the entire infection cycle. TayeBlu’s adsorption rate constant and total extent of adsorption were both lower by an order of magnitude in minimal medium than in rich medium (Fig. **2**A). Ammonium amendment of the minimal medium substantially rescued adsorption, although it remained significantly slower than in rich medium (*p* = 7.7 *×* 10^-12^, Welch modified two-sample t-test). Similarly, bursts were both later and far smaller (*p* = 5.364 *×* 10^-6^, Welch modified two-sample t-test) in minimal medium than in rich medium (Fig. **2**B), and nitrogen amendment of minimal medium produced burst dynamics closer in timing and scale (though still significantly smaller; *p* = 0.00641) to those in rich medium.

**FIG 2.**
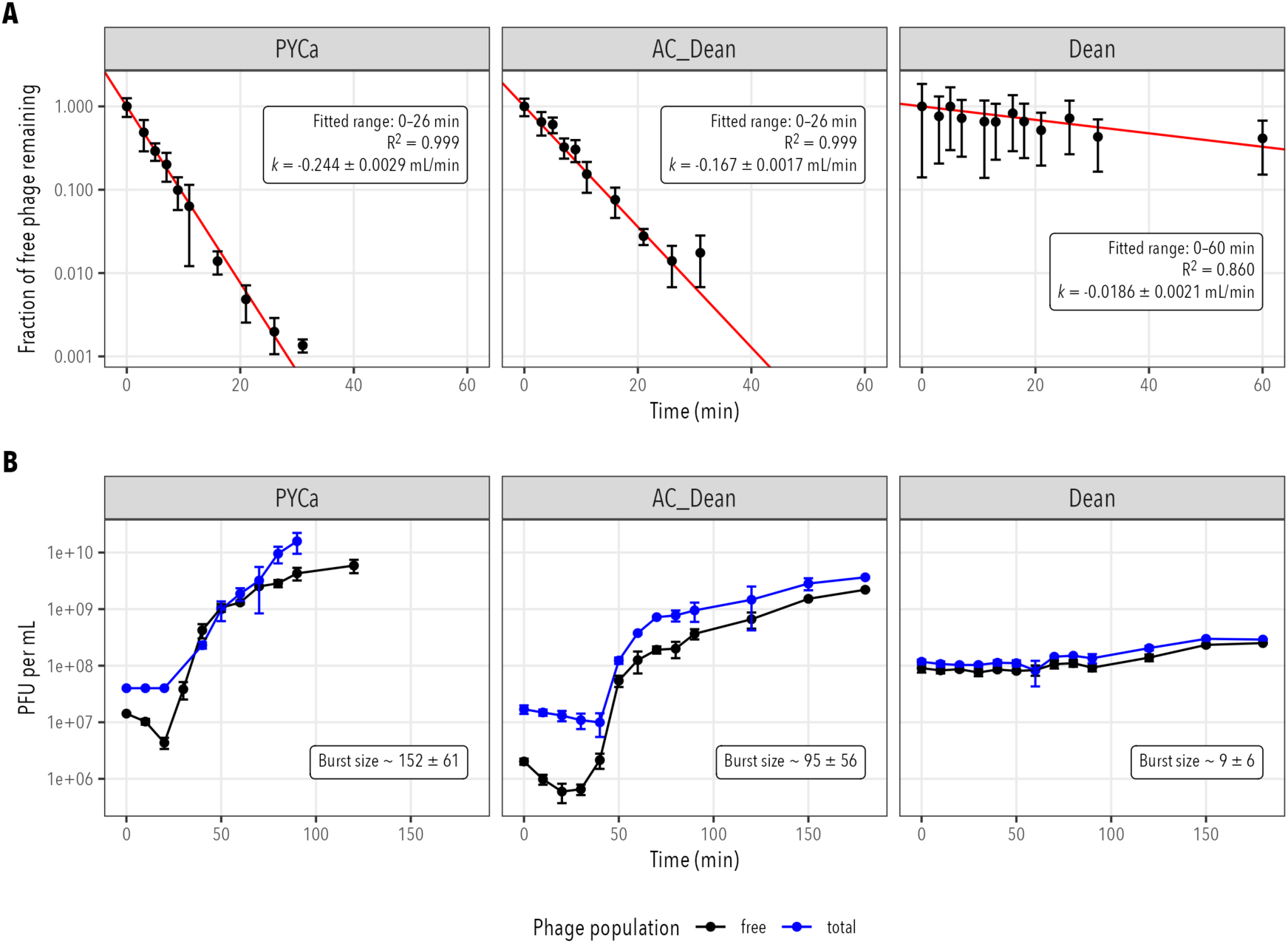
TayeBlu infection physiology on *A. vinelandii* strain OP, in rich (PYCa), N-replete minimal (AC_Dean), and minimal media (Dean). Each datapoint represents the mean of 3–6 independent measurements, weighted by Poisson error estimates; error bars show the standard deviation of the weighted mean. (A) Adsorption kinetics. Red lines show linear fits to natural log-transformed data in the stated range. Values for *k* are fitted slopes *±* standard error. (B) One-step infection curves. Burst size estimates (weighted mean *±* standard deviation) are based on the first three and last three timepoints for each medium (see Materials and Methods).

The extent of rescue by ammonium amendment (AC_Dean) of minimal medium strongly suggests that TayeBlu infection dynamics in minimal medium are shaped primarily by the requirement for diazotrophy in these conditions. Previous studies have shown that nutrient availability can modulate the expression of outer membrane proteins and lipopolysaccharides that commonly serve as phage receptors [84, 85]. In addition to any such changes, *Azotobacter* in well-aerated minimal medium is expected to grow diazotrophically and to deploy nitrogenase-protective mechanisms that are likely to alter phage infection. First, under N-fixing conditions, many *Azotobacter* strains substantially increase production of the extracellular product alginate, thought to limit diffusion of O_2_ from the medium to the cell [86]. Although *A. vinelandii* strain OP is generally considered a "non-gummy", alginate non-producing strain due to a spontaneous loss of function in its *algU* gene [87], we have frequently observed that a gummy phenotype spontaneously re-emerges during propagation under N limitation (Fig. S2); if present, an alginate capsule or other extracellular polysaccharide layer could easily alter receptor availability for phage, hampering adsorption [88]. Second, *Azotobacter* greatly increases its respiration rate to draw down intracellular O_2_, creating high C-substrate demand under N limitation [27, 89, 90]. In minimal medium, this high respiration rate could compromise resource availability for phage reproduction, extending the latent period and limiting burst size. The large change to infection dynamics observed here between rich and minimal media demonstrates the potential variability of TayeBlu impacts across soils with varying nutrient availability, highlighting the importance of characterizing soil phage infection dynamics across a range of growth conditions to support quantitative modeling of ecosystem impacts [91].

### Genome analysis

Complete genome sequencing revealed that TayeBlu possesses a circular double-stranded DNA genome of 59,885 bp with a G+C content of 50.34% (Table **1**). Structural and functional annotation using automated tools followed by manual curation in Apollo predicted 100 coding sequences (CDSs) with no identifiable tRNA genes. We identified 15–18 putative transcription terminators from the 30 predicted using ARNold [71] (Table **1**, Table S1). Start codon usage showed a predominance of ATG, with lesser usage of GTG and TTG. Functional annotation of the genome classified the CDSs into several categories (Fig. **3**): structural and assembly genes (21 genes, comprising head and packaging proteins, connector proteins, and tail proteins). DNA, RNA and nucleotide metabolism (16 genes), lysis (3 genes), transcription regulation (3 genes); other functions (11 genes), while the rest of the genes encode hypothetical proteins (45) with no known function (Fig. **3**).

**FIG 3.**
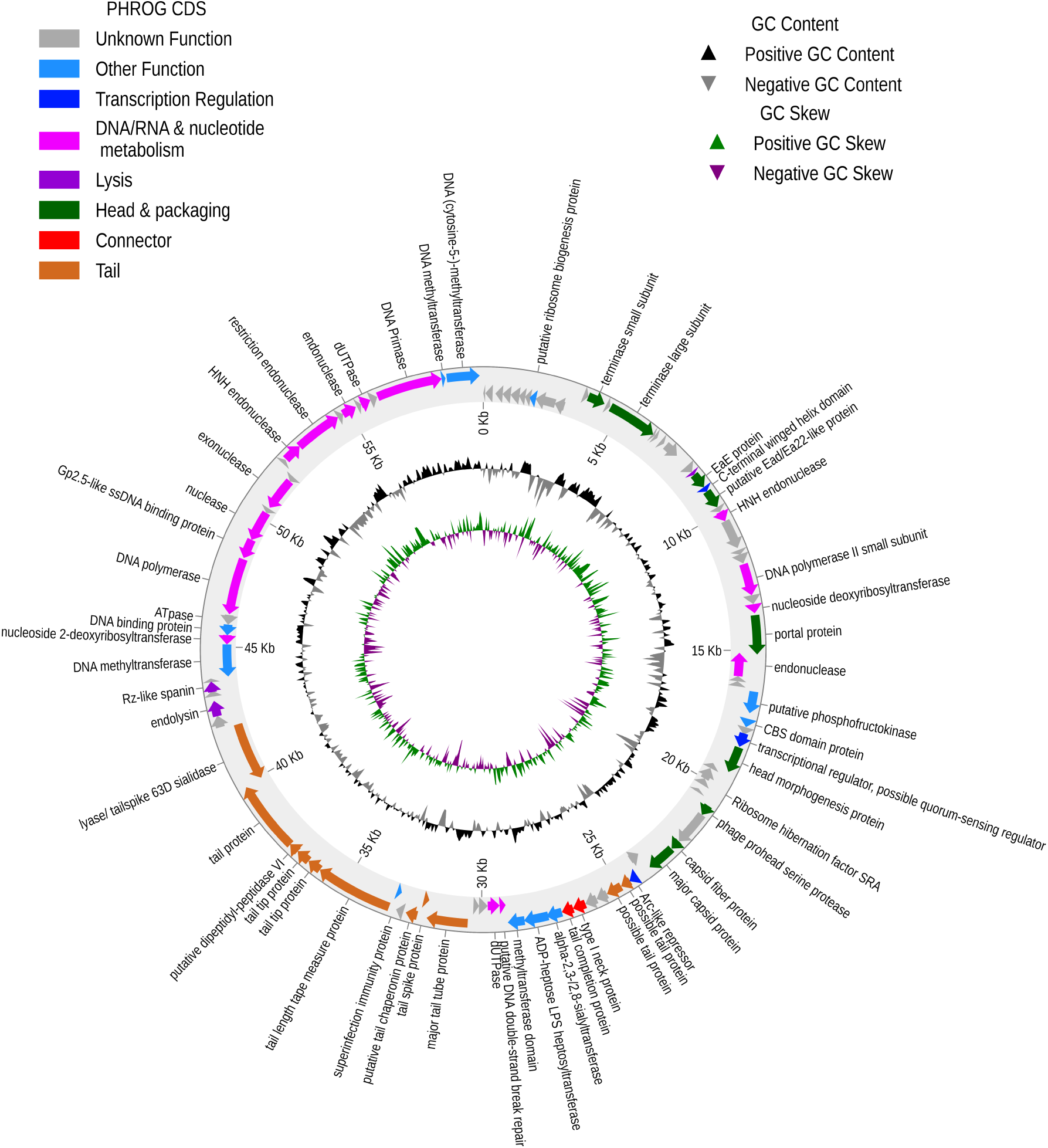
Complete circular genome of novel phage TayeBlu. Manually curated gene annotations (arrows, outermost rings) are colored by PHROG functional category. Nucleotide-level genome composition patterns are shown in the middle (GC content) and inner (GC skew) rings.

**TABLE 1.**
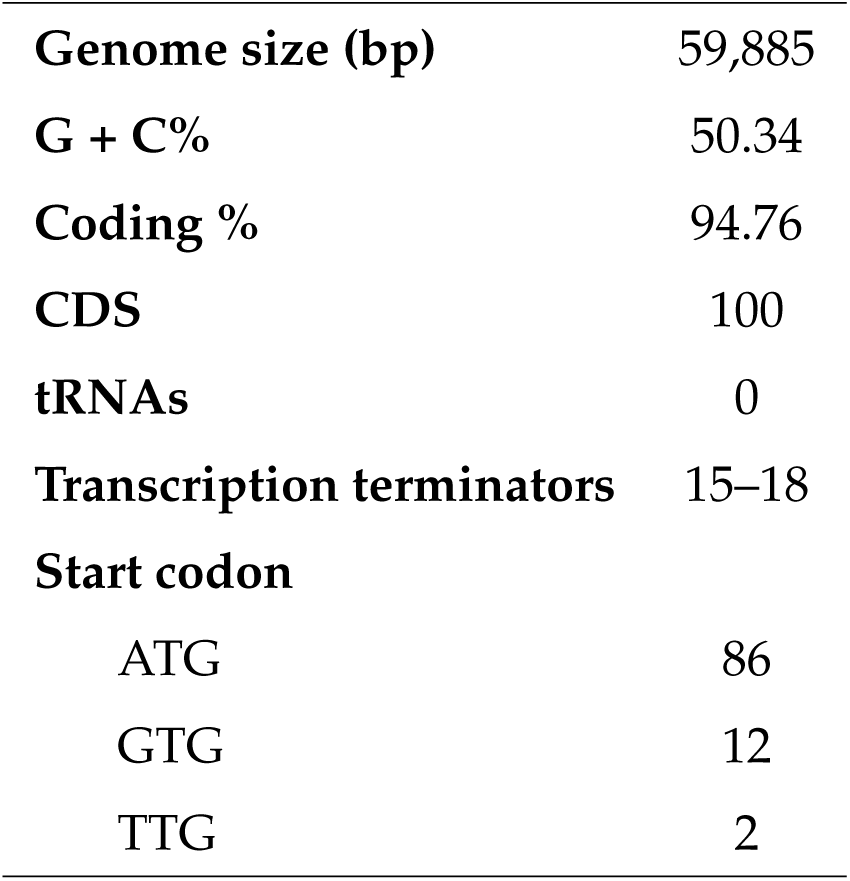
Genomic features of bacteriophage TayeBlu.

### Genes with potential functional significance

Intriguingly, TayeBlu encodes a putative phosphofructokinase (locus AZP_TayeBlu_36), previously reported as an AMG in marine and gut viruses [92, 93]. This enzyme catalyzes the conversion of fructose-6-phosphate to fructose-1,6-bisphosphate, an ATP-consuming early step in the Embden-Meyerhof-Parnas (EMP) glycolytic pathway that is the point at which a substrate glucose molecule is energetically committed to the pathway. Thus, any phage-mediated expression of phosphofructokinase during infection should partition glycolytic flux toward this pathway and away from the competing Entner-Doudoroff (ED) pathway. Fluxomic analysis of *A. vinelandii* has shown that only a small minority of glycolytic flux in uninfected cells uses EMP, while the bulk is directed through ED despite its lower ATP yield [89]. Hypothesizing that the tradeoff lowers the N demand for glycolytic enzyme synthesis, Wu *et al.* argue that *A. vinelandii* tunes central C metabolism so that the redox state of the cell supports respiratory protection of nitrogenase [89]. Phage-mediated rebalancing of glycolytic flux might thus put nitrogenase at risk. We hypothesize that TayeBlu will express phosphofructokinase during infection under conditions where phage replication is limited more by ATP than by N availability.

Beyond central C metabolism, several genes annotated in TayeBlu may influence phage-host interactions. Putative carbohydrate-active enzymes (CAZymes) include a lyase/tailspike with 63D sialidase homology (locus AZP_TayeBlu_74) and an alpha-2,3-/2,8-sialyltransferase (AZP_TayeBlu_57), which might interact with *A. vinelandii*’s exopolysaccharides, including alginate. Viral sialidases often facilitate cell entry or exit by modifying surface glycans [94]; earlier work has demonstrated alginate lyase activity in an unsequenced *A. vinelandii* phage [32]. Additionally, we identified a putative paratox (Prx) domain protein (AZP_TayeBlu_39) with sequence similarity to inhibitors of the quorum sensing receptor ComR [95]. Quorum sensing regulation plays a critical role in biofilm formation, e.g., by altering production of extracellular polymeric substances including alginates (reviewed in [96]); a paratox protein might allow TayeBlu to manipulate host cell signaling to alter biofilm formation or EPS production. Finally, the genome encodes a protein with homology to recently described ribosome hibernation factors (AZP_TayeBlu_43) [97], which may influence host translation machinery under nutrient-limited conditions. The roles of these genes in the TayeBlu infection cycle across growth conditions remain to be experimentally validated.

### Taxonomic placement

Following current ICTV guidelines [83], we sought to classify TayeBlu on the basis of its genome [98]. The proteome clustering tool vConTACT3 [74] placed TayeBlu in established taxonomic ranks to class level, in realm *Duplodnaviria*, phylum *Uroviricota*, and class *Caudoviricetes*, consistent with its identification as a tailed bacteriophage [98]. At lower taxonomic levels, TayeBlu was assigned to novel_order_46, novel_family_19, novel_subfamily_1, and novel_genus_0 (Fig. S3), indicating it represents a previously uncharacterized lineage within *Caudoviricetes*. TayeBlu clustered with 5,056 *Caudoviricetes* viruses in Viral RefSeq, with 510 members classified under novel_order_46. Within novel_family_19, only five other viruses were identified: Klebsiella phage vB_KpnS-Carvaje (56,858 bp, 97 proteins), Proteus phage VB_PmiS-Isfahan (54,836 bp, 96 proteins), Salmonella phage 9NA (52,869 bp, 80 proteins), and Salmonella phage vB_SenS_Sasha (53,263 bp, 80 proteins). At the subfamily and genus levels, TayeBlu is the sole representative of novel_subfamily_1 and novel_genus_0, mapping new territory in known viral diversity.

To assess the phylogenetic placement of TayeBlu within the still-undescribed novel_family_19, we generated a phylogenetic tree using all proteomes in VipTree S4 based on proteome-wide similarities. The analysis revealed that TayeBlu forms a single, cohesive monophyletic lineage with the four vConTACT3-assigned family members (Fig. **4**). This confirms that they share a significant number of orthologous genes, consistent with ICTV family-level classification standards. Next, we scored protein similarity to phages in the Virus-Host DB [99], identifying a total of 42 phages with moderate similarity (ViPTree similarity score *≥* 0.020) to TayeBlu, including the four vConTACT3-identified members of novel_family_19. Two of these 42 shared a node in the ViPTree with TayeBlu and the four vConTACT3 relatives.

**FIG 4.**
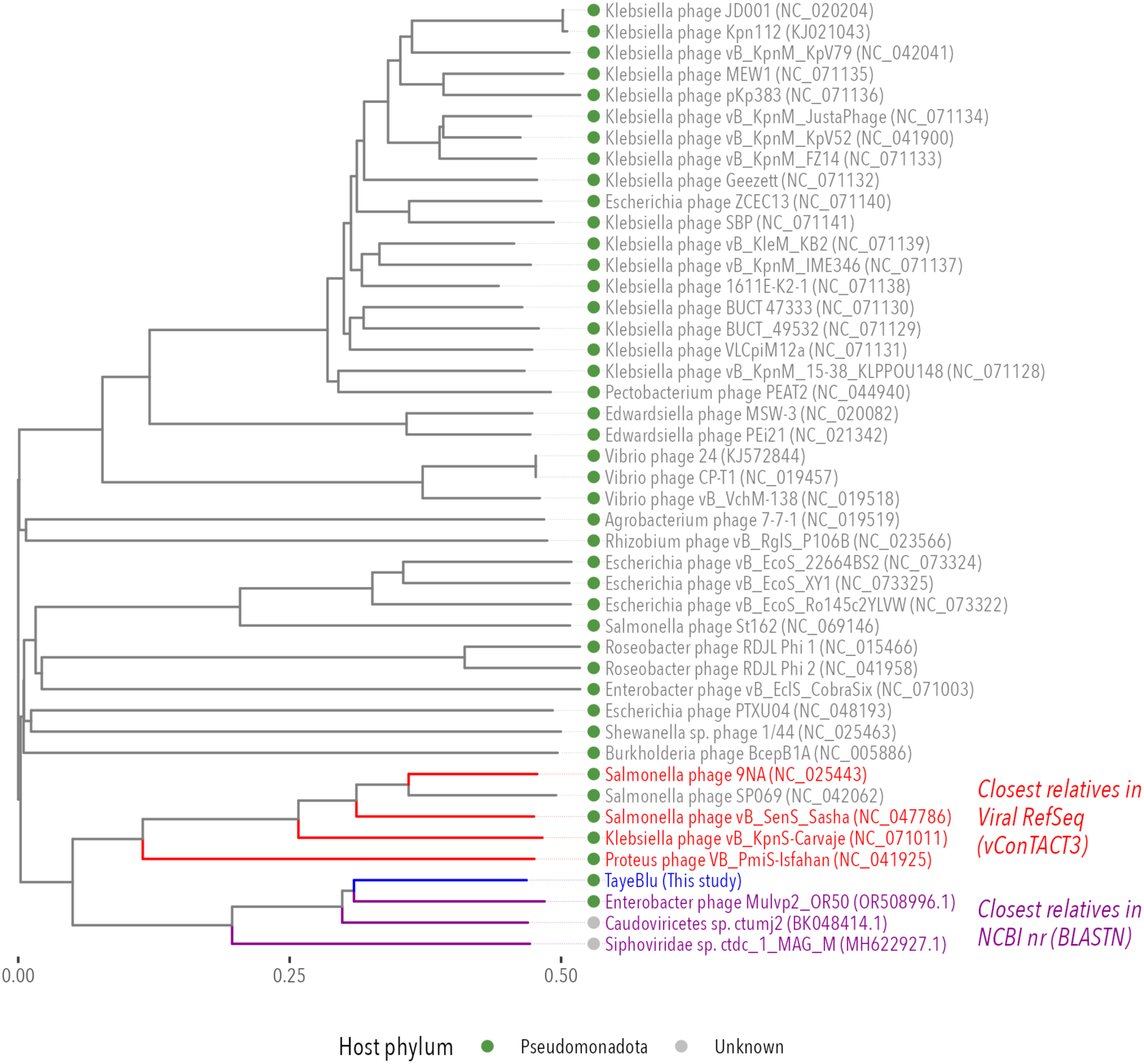
Proteome-based phylogenetic relationships between TayeBlu (blue), three close relatives identified in NCBI nr by genomic similarity (purple), and 41 additional relatives with ViPtree proteome similarity scores of at least 0.020 (gray), including the four members of novel_family_19 identified by vConTACT3 (red). Nearly all viruses in the tree are known to infect hosts from phylum Pseudomonadota (green dots); the remaining two phage genomes were recovered from metagenomic sequencing and their hosts are unknown (gray).

Finally, to identify additional relatives of TayeBlu, we performed BLAST searches against all viruses in the NCBI Nucleotide collection (nr/nt) database against the complete TayeBlu genome. The search revealed no highly similar relative but three moderately similar phages with nontrivial query coverage: Enterobacter phage Mulvp2 (OR508996.1; 67% query coverage, 93.93% identity), *Caudoviricetes* sp. isolate ctumj2 (BK048414.1; 64% query coverage, 93.78% identity), and Siphoviridae sp. ctdc_1 (MH622927.1; 26% query coverage, 72.38% identity). In this expanded group, all phages whose hosts are known infect hosts within the phylum *Pseudomonadota* (Fig. **4**).

When we repeated vConTACT3 protein clustering with the close ViPTree and BLASTN relatives included, novel_family_19 expanded to include not only TayeBlu and the four previously identified RefSeq members but also the three phage identified by BLASTN and one of the two phage identified by ViPTree (Table S2). (The remaining close ViPTree relative was identified as a much shorter satellite phage and excluded from further analysis.) This cluster structure indicates that protein-level homology supports a shared evolutionary origin for these nine phages. ViPtree proteome phylogeny grouped TayeBlu and the three BLASTN hits with the close ViPTree and vConTACT3 hits but in a distinct and cohesive sister subclade (Fig. **4**).

Analysis of novel_family_19 at the nucleotide level suggests substantial diversification both between and within the two sub-clades evident in the proteome tree. As expected, VIRIDIC intergenomic similarity analysis showed that the closest relative to TayeBlu was its top BLAST hit, sharing only 65.3% ANI (Fig. **5**). Notably, TayeBlu exhibited minimal nucleotide similarity (≤10%) with the ViPTree and vConTACT3 family members in the second subclade. This observation is consistent with the growing recognition that phage taxonomy should rely primarily on protein conservation patterns, as the extent of diversification among related phage can obscure relationships at the nucleotide level [100, 101] and, further, that family-level groupings represent broader genetic diversity among bacteriophage than among eukaryotic viruses [102]. With current ICTV intergenomic similarity thresholds of *≥* 95% for the species level and *≥* 70% for the genus level, the highest intergenomic similarity observed here (67.6%) falls well short of the genus-level threshold. Our findings therefore suggest that each identified member of this novel family represents a distinct genus and species within this viral lineage.

**FIG 5.**
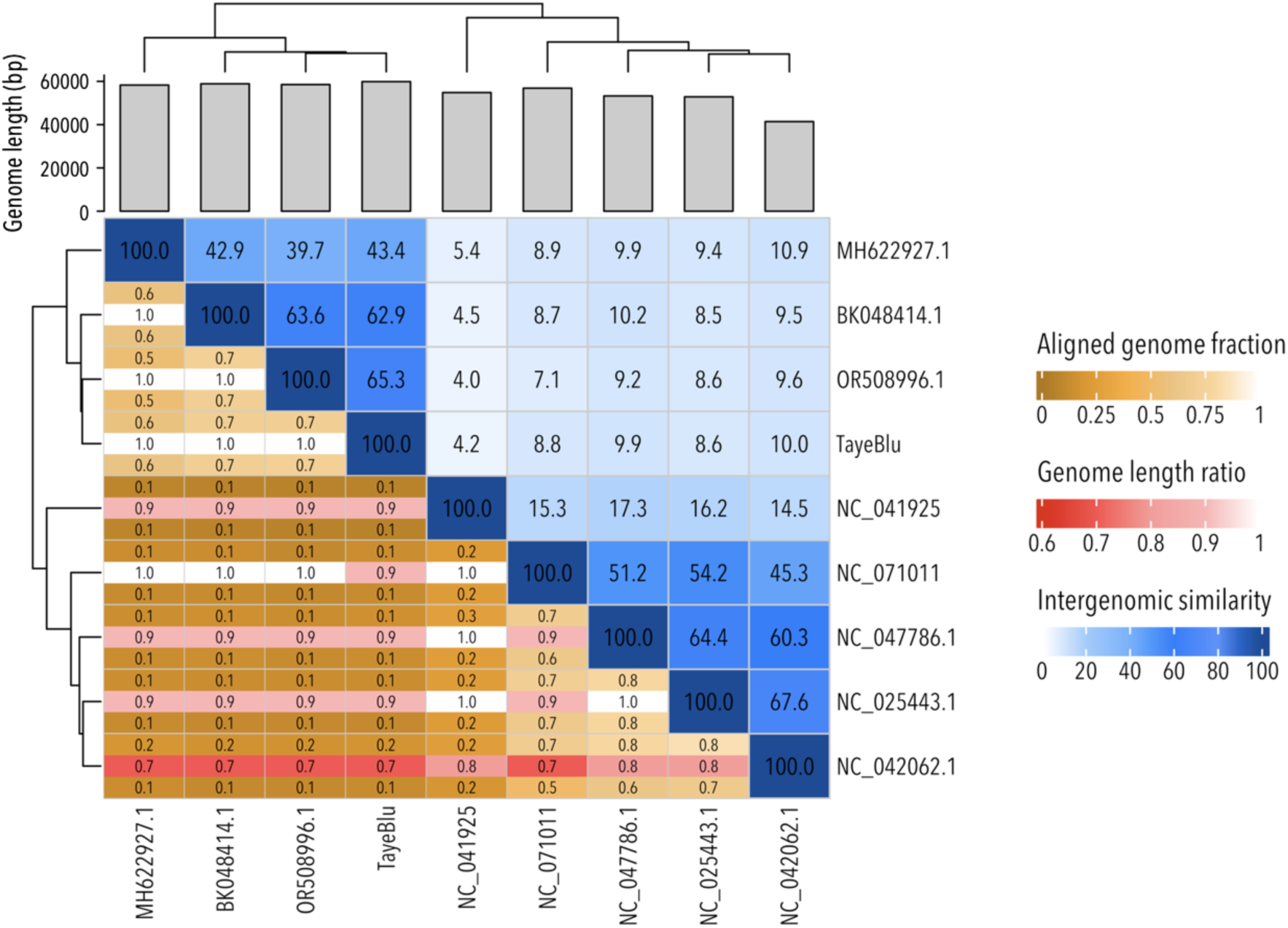
VIRIDIC intergenomic similarity matrix for TayeBlu and the other identified members of the novel family highlighted in this study.

### Genome comparison and core gene identification

TayeBlu and its eight identified relatives have genomes of 41–59 kb with 69–100 predicted ORFs (Fig. **6**), a total of 799 ORFs across the family. We applied protein clustering (sequence identity *≥* 30%, coverage *≥* 80%) to identify this novel family’s core genes, finding 397 unique clusters (Table S3, Fig. S5 of which 11 were present in all nine examined genomes. These 11 core genes, ∼12% of the total open reading frames (ORFs) in this phage family, constitute the family’s conserved genetic framework. An additional five genes were shared between TayeBlu and all but one phage in the family, extending the near-core genome to 16 genes. Functional annotation of these conserved genes revealed that they primarily encode structural components (capsid and tail proteins), DNA packaging machinery (terminase and portal proteins), and enzymes involved in DNA replication (DNA helicase, polymerase, dUTPase), alongside an endonuclease. Genome regions encoding structural tail proteins and DNA packaging machinery also show substantial synteny (Fig. **6**).

**FIG 6.**
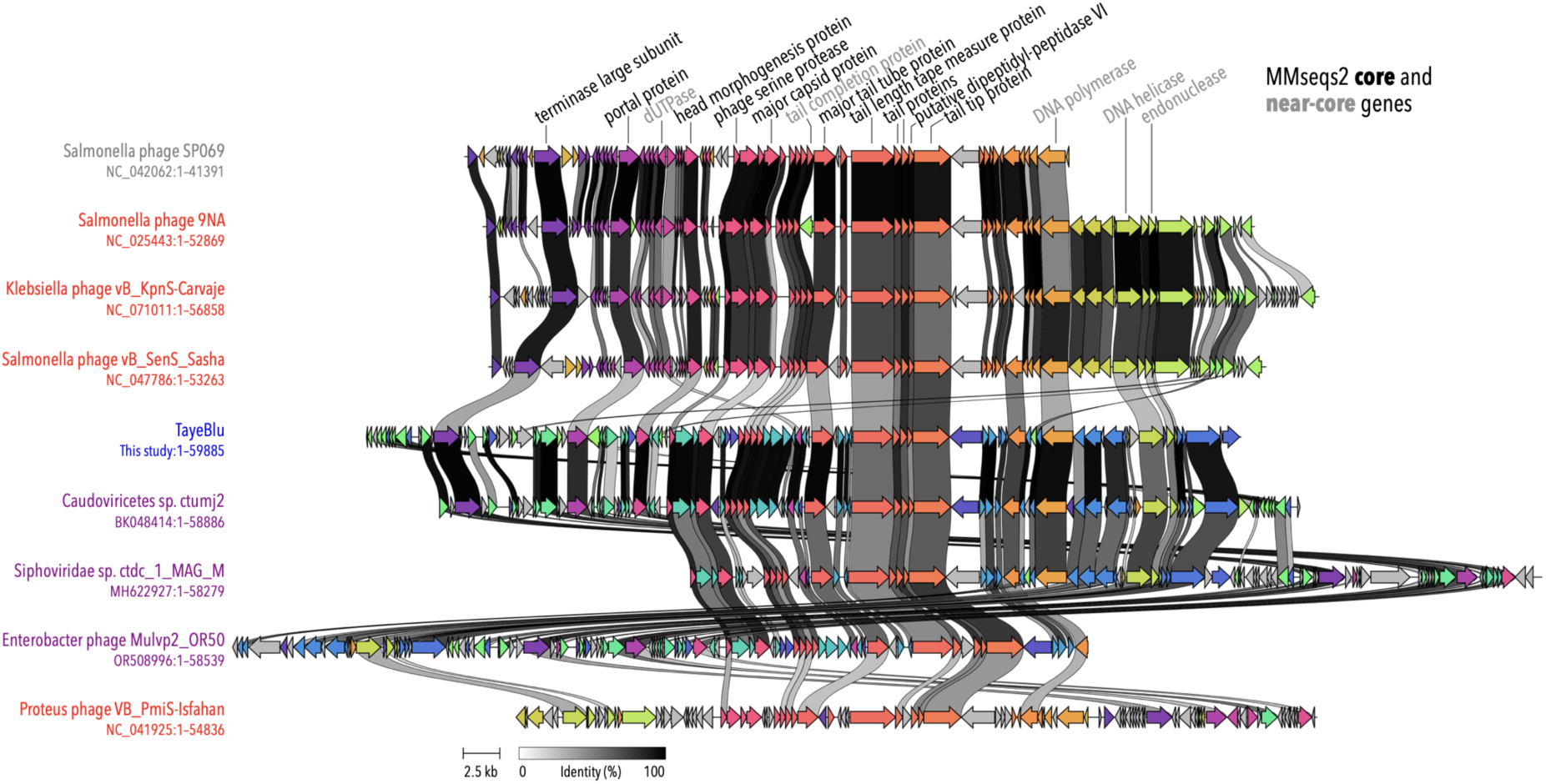
Core genes and synteny relationships in novel_family_19. Sequence names are colored as in Fig. **4**: blue, TayeBlu; purple, phage identified by BLASTN in NCBI nr/nt; red, identified by vConTACT3 in Viral RefSeq; gray, identified by ViPTree in the Virus-Host DB. Ribbons connect shared genes identified by Clinker alignments; ribbon shading is scaled to pairwise percent identity at the protein level. Core genes (gene functions labeled in black) were identified by MMseqs2 protein clustering as present in all phage and near-core genes (gray) as present in 8 of 9 phage.

The predominance of structural and assembly genes in the core genome is consistent with previous observations that these functional categories tend to be the most highly conserved genes across diverse phage lineages [103, 104]. The critical importance of these functions in the phage life cycle likely places these genes under higher selective pressure than the rest of the genome, as seen previously in tailed bacteriophage large terminase genes (reviewed in [105]). The limited but syntenic core genome suggests that the essential framework for virion assembly and DNA replication has remained intact during adaptive radiation to the family’s ecological niches. Notably absent from the core genome are genes involved in transcriptional regulation and host takeover functions (Fig. S5), suggesting diverse host interaction strategies. The conservation pattern observed in this novel phage family, with structural proteins, DNA packaging components, and replication machinery forming the stable core within a family marked by low overall genome similarity, is consistent with the observation that phages can maintain a stable core genome over extended periods while acquiring variable complements of genes through recombination from the phage pan-genome [106], highlighting the balance between functional conservation and adaptive diversification.

### Clustering with phages isolated from natural soil and rhizosphere

Finally, we investigated TayeBlu’s genomic relatedness to other soil and rhizosphere-associated phages. We clustered TayeBlu’s genome against 103,626 soil and rhizosphere viral operational taxonomic units (vOTUs) from the PIGEON database [82]. We found no closely related phages within this extensive soil-associated dataset, suggesting that TayeBlu represents a novel viral lineage. Soil viromes are known to harbor extensive genetic novelty [107, 108], and this vast diversity is still largely unexplored [109]. Even across different rhizosphere habitats, phages display exceptionally diverse communities depending on plant species, soil type, and geographic location [110, 109, 111], with most identified viral sequences showing little similarity to known phages from any environment [110]. This suggests highly specialized adaptation to ecological niches that are subject to plant-soil-microbe interactions. The lack of phage genomes closely related to TayeBlu underscores the limited representation of soil viral diversity in current databases and the pressing need for expanded efforts to catalog and study phages from diverse terrestrial ecosystems.

### Conclusion

The novel phage TayeBlu, isolated on *Azotobacter vinelandii* strain OP from agricultural rhizosphere soil, is a member of a novel family and represents a distinct genus and species within this viral lineage. Its infection physiology varies dramatically depending on nutrient availability and particularly on the exogenous supply of fixed nitrogen to its facultatively diazotrophic host. We hypothesize that this variation will alter the phage’s ecological impacts across different soils and agricultural practices, potentially influencing bacterial community structure and evolution [112, 113], N flow within the rhizosphere, and ecosystem outputs from the globally significant process of biological N fixation. Genomic analysis reveals extensive diversification even among the few members of this phage family, highlighting the need for future isolation efforts to illuminate the ecological roles and impacts of phage in soil systems.

## ACKNOWLEDGMENTS

We thank Robert Ward, Marissa Gittrich, Natalie Solonenko, Marie Burris, members of the Bagby lab for useful discussions. We thank Jessica LaBella, Isabella Darling, and Madison Talley for technical assistance with one-step experiments. We gratefully acknowledge an award from the Case Western Reserve University Department of Biology Oglebay Fund to TMA and CWRU SOURCE summer funding to ML.

## DATA AVAILABILITY STATEMENT

The complete genome sequence of phage TayeBlu has been submitted to GenBank (submission #2951289, accession number pending NCBI review).

## Supplemental material

Supplementary Figures S1–S5

Supplementary Table S1

Supplementary Table S2

Supplementary Table S3

