## Supplemental Figures 1-5 for "Isolation and characterization of TayeBlu, a novel bacteriophage of *Azotobacter vinelandii*"

### Supplementary Figures

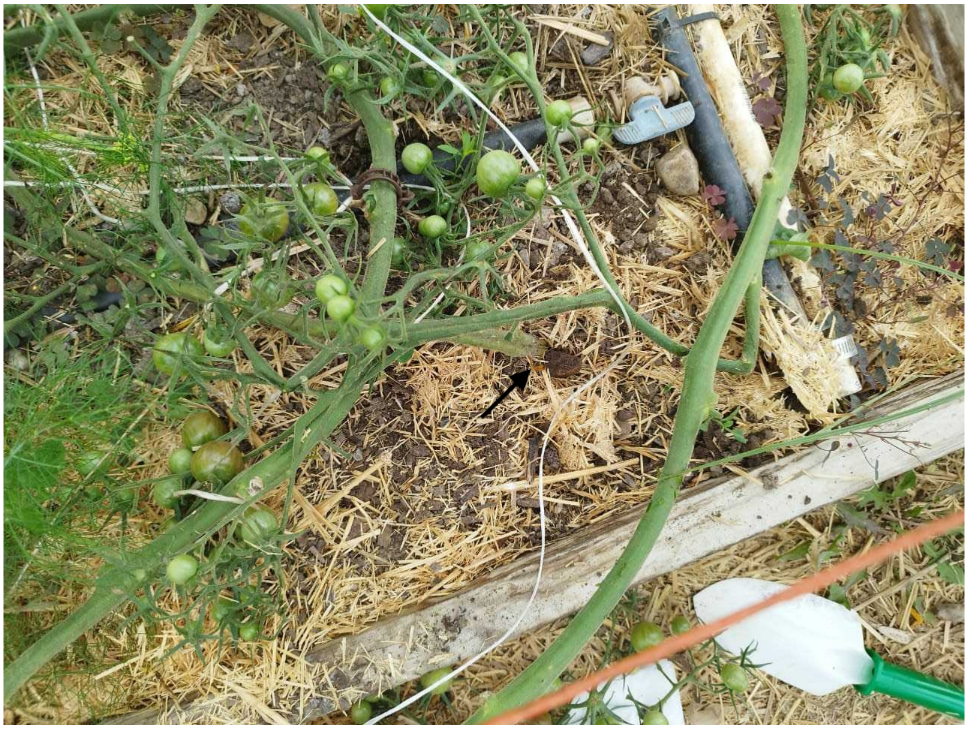

**FIG** S1 Tomato plant at the Case Western Reserve University Farm greenhouse prior to rhizospheric soil collection. Soil was sampled from the root-adjacent zone at the plant base (black arrow) using a sterilized hand shovel, targeting areas with visible fine roots.

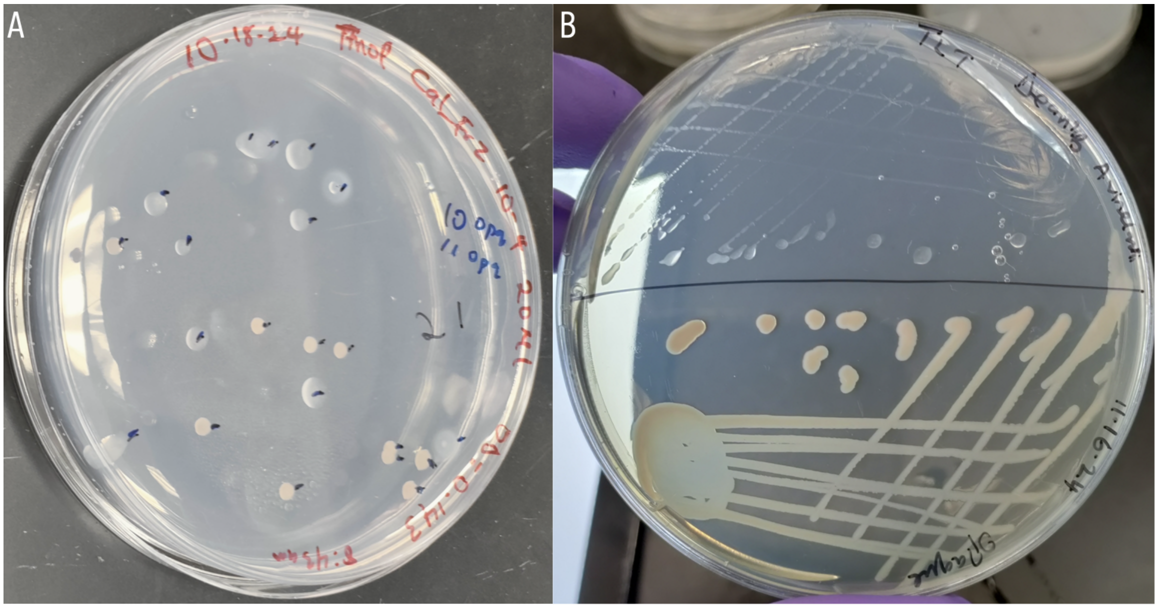


**FIG** S2  Phenotypic variation in *Azotobacter vinelandii* strain OP culture after growth in Dean's B liquid medium. (A) Spread plate of *A. vinelandii* strain OP culture on Dean's B medium at $10^{-4}$ dilution showing two distinct colony morphologies: gummy (larger, translucent, pearlescent) and non-gummy (compact, opaque, greenish-white) variants. (B) Streak plate on Dean's B medium demonstrating the isolated gummy (top) and non-gummy (bottom) morphotypes.

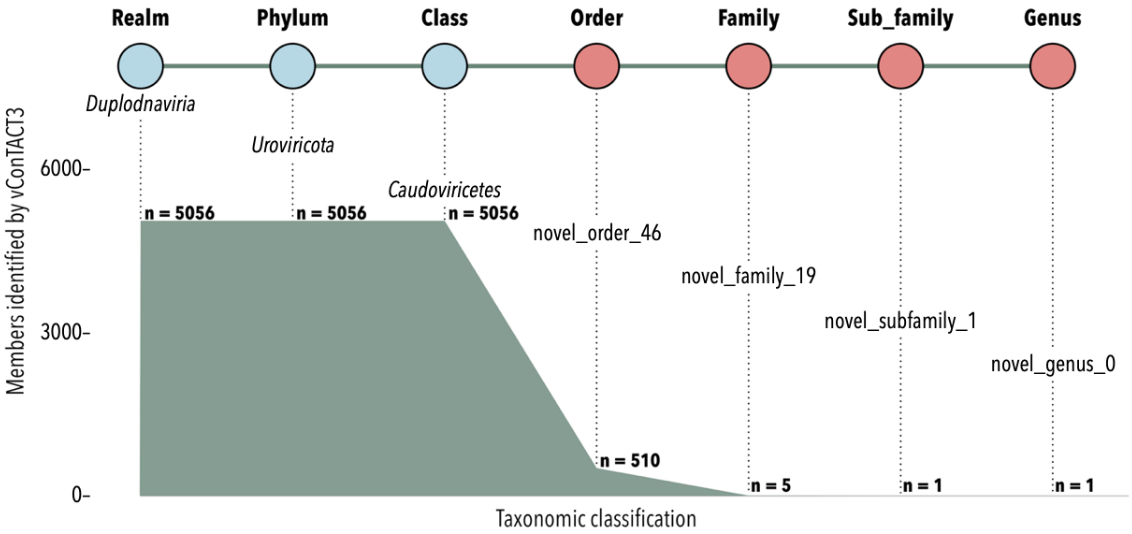


**FIG** S3 Taxonomic classification of TayeBlu by vConTACT3, showing counts of phage in Viral RefSeq that share TayeBlu's taxonomy at each level. Counts include TayeBlu.

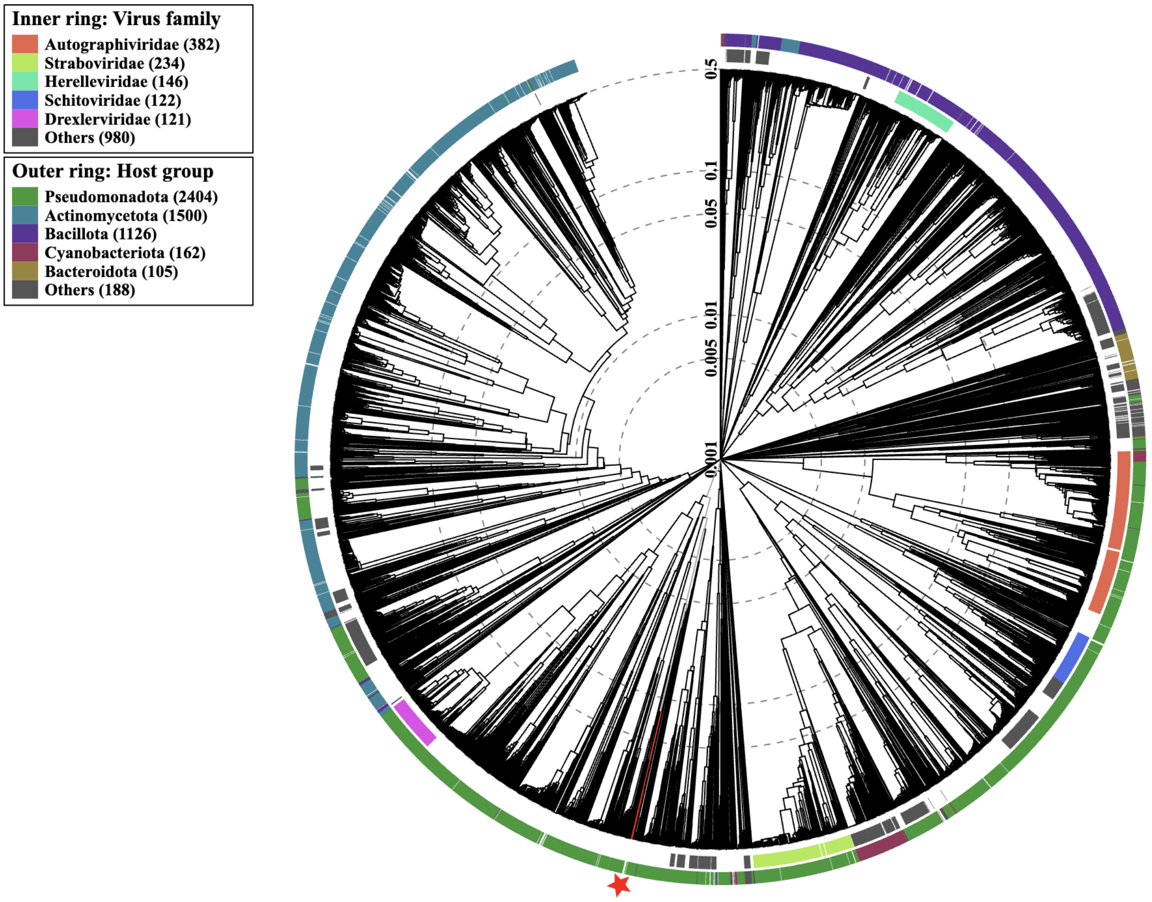


**FIG** S4 Protein-based phylogeny of all TayeBlu relatives identified by ViPTree in Virus-Host DB, showing viral families (inner ring) and host group (outer ring). TayeBlu is marked by a red star. TayeBlu and the four phage identified by vContACT3 are indicated by red branches.

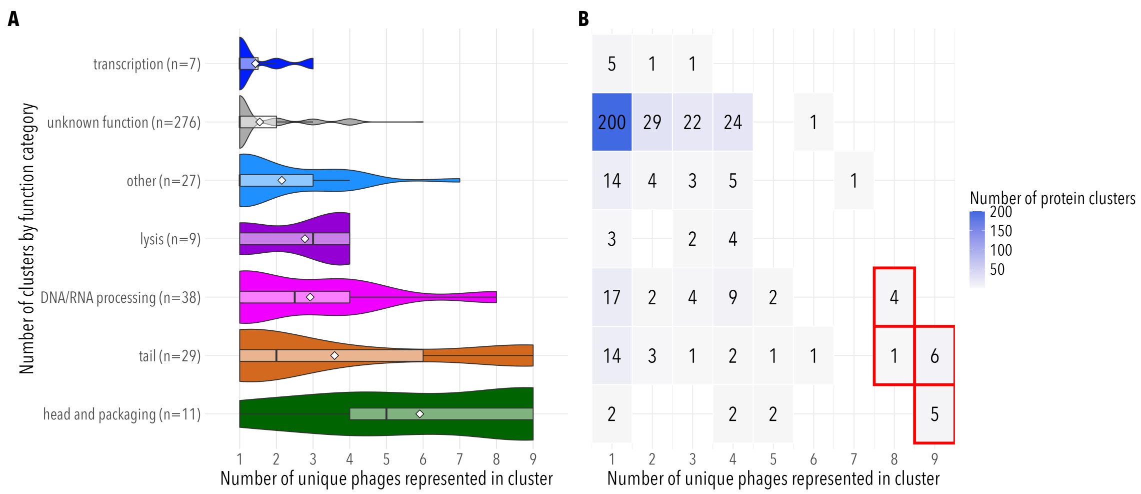


6

**FIG** S5 Distribution of protein cluster membership across the phage family by functional category. (A) Boxplot and violin plot. Mean, white diamonds; median, crossbar; lower bound, 25th percentile; upper bound, 75th percentile. Functional categories are ordered by mean number of phages represented in per cluster. (B) Heatmap of cluster counts. Data is the same as in (A). Clusters boxed in red contain core and near-core genes.
